# SIV clearance from neonatal macaques following transient CCR5 depletion

**DOI:** 10.1101/2023.05.01.533682

**Authors:** Jesse D. Deere, David Merriam, Kawthar Machmach Leggat, Wen-Lan William Chang, Gema Méndez-Lagares, Hung Kieu, Joseph Dutra, Justin Fontaine, Wenze Lu, Ning Chin, Connie Chen, Bryant Chi-Thien Tran, Jessica Salinas, Corey N. Miller, Steven G. Deeks, Jeffrey D. Lifson, Kathleen Engelman, Diogo Magnani, Keith Reimann, Mario Stevenson, Dennis J. Hartigan-O’Connor

**Affiliations:** Department of Medical Microbiology and Immunology, School of Medicine, University of California Davis, Davis, California, USA; California National Primate Research Center, University of California Davis, Davis, CA, USA; Walter Reed Army Institute of Research, Silver Spring, MD USA; Department of Medicine, University of Miami, Miller School of Medicine, Miami, Florida, USA; Division of Infectious Diseases, University of Miami, Miller School of Medicine, Miami, Florida, USA; Diabetes Center, University of California San Francisco, San Francisco, California, USA; Department of Medicine, University of California San Francisco, San Francisco, California, USA; Department of Medicine, University of California San Francisco, San Francisco, CA, USA; AIDS and Cancer Virus Program, Frederick National Laboratory for Cancer Research, Frederick, MD, USA; Nonhuman Primate Reagent Resource, MassBiologics of the University of Massachusetts Medical School, Boston, Massachusetts, USA; Division of Experimental Medicine, Department of Medicine, University of California San Francisco, San Francisco, CA, USA

## Abstract

Treatment of people with HIV (PWH) with antiretroviral therapy (ART) results in sustained suppression of viremia, but HIV persists indefinitely as integrated provirus in CD4-expressing cells. Intact persistent provirus, the “rebound competent viral reservoir” (RCVR), is the primary obstacle to achieving a cure. Most variants of HIV enter CD4^+^ T cells by binding to the chemokine receptor, CCR5. The RCVR has been successfully depleted only in a handful of PWH following cytotoxic chemotherapy and bone marrow transplantation from donors with a mutation in *CCR5*. Here we show that long-term SIV remission and apparent cure can be achieved for infant macaques via targeted depletion of potential reservoir cells that express CCR5. Neonatal rhesus macaques were infected with virulent SIVmac251, then treated with ART beginning one week after infection, followed by treatment with either a CCR5/CD3-bispecific or a CD4-specific antibody, both of which depleted target cells and increased the rate of plasma viremia decrease. Upon subsequent cessation of ART, three of seven animals treated with CCR5/CD3-bispecific antibody rebounded quickly and two rebounded 3 or 6 months later. Remarkably, the other two animals remained aviremic and efforts to detect replication-competent virus were unsuccessful. Our results show that bispecific antibody treatment can achieve meaningful SIV reservoir depletion and suggest that functional HIV cure might be achievable for recently infected individuals having a restricted reservoir.

## MAIN TEXT

Instances of HIV cure or remission have been rare. Notable cases of apparent HIV cure have involved PWH who were treated aggressively for underlying malignancies with regimens that included radiation and/or cytotoxic chemotherapy or immunotherapy and bone marrow transplantation from donors with a mutation in chemokine receptor 5 (CCR5Δ32/Δ32), known to provide resistance to HIV replication.^1^ The results suggest that elimination of cells bearing this chemokine receptor can make an important contribution to eradication of the virus.

Instances of complete elimination of virus (a “cure”) in the non-human primate model have been similarly rare. ART is not curative in either humans or macaques once a reservoir has been fully established, within approximately three days after infection. No instance of apparent cure was seen when ART was started three or seven days after SIV or SHIV infection (0/8 and 0/11 in refs. 8 and 9, respectively). In another example, 39 animals were infected intravenously with two focus-forming units of SIVmac239X^10^, an inoculum that is both clinically relevant and predicted to result in slow spread of the virus, and were treated from 4-12 days after infection. In this study, as well, no cure was demonstrated when ART was started seven days after infection (0/10) and only one of 13 animals treated six days after infection failed to rebound (1/13). Animals that started ART 4 or 5 days post-infection universally rebounded.

Other approaches to achieving post-ART control have been devised, most of which combine strategies for stopping viral replication and eliminating or controlling the RCVR.^11^ When viral replication is controlled by ART, reservoir reduction might be approached via gene editing; latency reversal combined with immunologic targeting of re-activated cells; “block-and-lock”;^12^ or direct depletion, e.g., targeting of surface markers known to be expressed on infected cells and/or cells vulnerable to infection. One of the most successful approaches to date combined the broadly neutralizing antibody PGT121 with the TLR7 agonist vesatolimod, leading to apparent cure in 5/11 adult animals previously treated with ART beginning seven days after SHIV infection.^9^ Direct depletion of the RCVR has received comparatively little attention. The approach is counter-intuitive because, given the lack of a specific marker for latently infected cells, a large number of uninfected lymphocytes must be depleted with the longer-term goal of preserving immune function. Nonetheless, elimination of host immune cells, presumably encompassing the RCVR, was the first step in both instances of human HIV cure; furthermore, it is possible that depletion of Env-expressing cells via ADCC made a contribution to the success of studies employing PGT121. Therefore, we hypothesized that depletion of infected and infectable cells could permit SIV cure if accomplished soon after infection when the RCVR remains limited in extent and uniform in terms of associated surface markers. We predicted that such treatment in infants would be followed quickly by robust, long-term T cell reconstitution due to active thymic output.

To test this hypothesis, we selected two agents predicted to eliminate both SIV-infected cells contributing to the RCVR and uninfected potential target cells: an anti-CD4 antibody and an anti-CD3/CCR5 bispecific antibody (bsAb). The bsAb is intended to juxtapose CCR5-expressing cells with cytotoxic effectors. The infection and treatment protocol modeled perinatal infection followed by early ART, with administration of the experimental depleting agent soon thereafter (Fig. 1A). To maximize signal for effects that might prove clinically meaningful, we chose high-dose oral infection with virulent SIVmac251. Neonatal macaques were infected at two weeks of age by oral inoculation with 50,000 TCID_50_ of SIVmac251, given twice on the same day. Antiretroviral “triple therapy” (tenofovir, emtricitabine, and dolutegravir^13^) was started one week later to permit time for peripheral dissemination of virus in all infected animals. Two days after ART initiation, two groups were administered candidate RCVR-depleting therapies: group B received the depleting anti-CD4 antibody, CD4R1, while group C received an anti-CD3/CCR5 bsAb previously described by our group to deplete CCR5^+^ cells.^14^ ART was withdrawn after 18 weeks and viral rebound was subsequently monitored.

**Figure 1.**
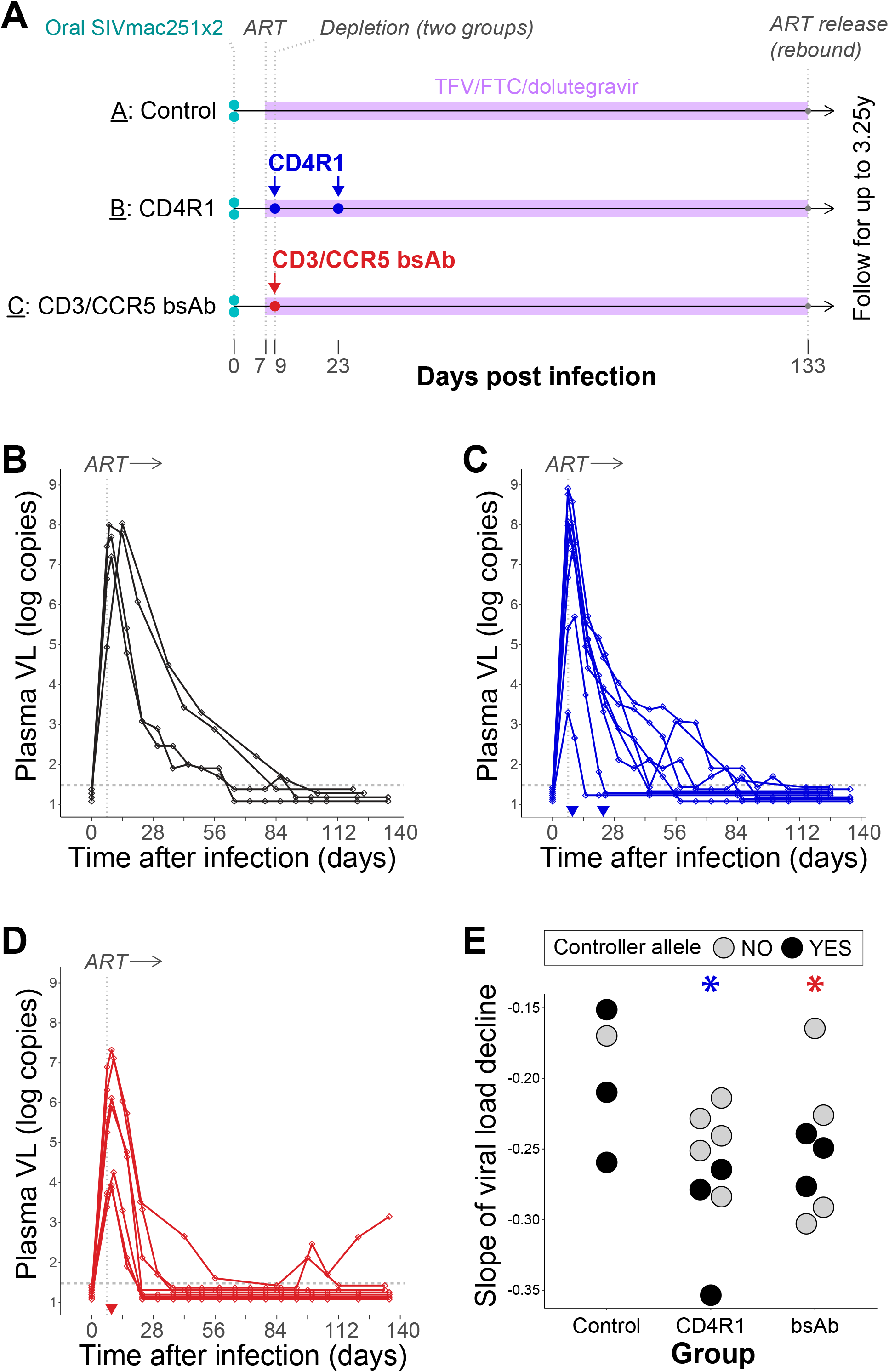
SIV infection of newborn macaques, early treatment, and depletion of potential reservoir cells. **(A)** Experimental schema. To model perinatal infection, newborn macaques were infected by oral inoculation (twice in one day) at two weeks of age. Antiretroviral therapy with tenofovir, emtricitabine, and dolutegravir was started seven days after infection. Depleting antibodies were given 2 and 16 days (group B; CD4R1) or only 2 days (group C, anti-CD3/CCR5 bsAb) after ART initiation. Therapy was withdrawn 19 weeks (133 days) after infection. **(B-D)** Plasma SIV RNA viral loads in the infected macaques of groups A, B, and C, respectively, before treatment interruption. **(D)** The slope of viral-load decline in each animal was calculated by linear regression using the peak value and those from the subsequent three weeks (asterisks indicate p<0.05 for contrast of this group with the control group).

19/24 challenged infants were infected by the oral inocula, with 3/5 uninfected infants having inherited a Mamu allele associated with post-acquisition control of viral replication (one A*01^+^/B*17^+^, one B*17^+^, one A*01^+^). Compared to those treated with ART alone, animals in groups B and C both experienced more rapid viral-load decay following antibody treatment (Fig. 1B-E; p=0.012 and 0.039 for slope of decay in groups B and C), suggesting either a rapid effect of depletion or a secondary effect on SIV spread, such as blockade of viral entry.^15^ Mamu allele inheritance did not predict viralload decay or time to achieve undetectable viral load. However, failure to inherit a controller allele was associated with viral “blips” after suppression was first achieved (3/9 animals with controller alleles and 9/10 without blipped; p=0.02 by Fisher’s test). Together, these data suggest that targeting CD4-or CCR5-expressing cells in acute infection accelerates decay of viremia.

### Long-lasting effect of bispecific antibody on blood and tissue immunophenotypes

Animals in group B exhibited a profound depletion of CD4^+^ cells from blood, including ∼100% of CD4^+^ monocytes (Extended Data Fig. 1) and ∼100% of CD4^+^ T cells (Fig. 2A). This depletion was durable, with a substantial deficiency of such cells persisting in 2/5 animals until 12 weeks after the first antibody administration (Fig. 2A). As expected, CD8^+^ T cells were not depleted (Fig. 2B). Animals in group C (bsAb recipients) exhibited temporary CD3^+^ lymphopenia (Fig. 2C-D) and specific, durable loss of CCR5^+^ cells in agreement with the pattern we previously described (Fig. 2E and ref. 14). Loss of circulating CCR5^+^ cells from blood was essentially complete, with <=3 CCR5^+^ CD4^+^ memory and <=5 CCR5^+^ CD8^+^ memory T cells detected in all treated animals for at least 14 weeks post bsAb administration (Fig. 2E, right panels showing absolute counts per ml of blood; evaluated using a non-cross blocking mAb to CCR5). Using the same bsAb, we previously demonstrated depletion of CCR5^+^ cells from lymph node and colon,^14^ so the depletion we observed in blood was not caused by trafficking of CCR5^+^ cells to those tissues. Surprisingly, relative depletion of circulating CCR5^+^ cells persisted in some treated animals for ≥34 weeks following a single administration of bsAb, while one macaque saw restored CCR5 expression by the same time point. We hypothesized that the idiopathic, transient CD3^+^ lymphopenia seen most prominently one week after bsAb administration might be accompanied by immune activation; however, we did not detect any statistically significant differences in inflammatory cytokine production between bsAb-treated animals and controls by multiplex Luminex assay (Extended Data Fig. 2; depletion of cells from tissues is addressed in ref. 14). Therefore, the RCVR-depleting agents tested here had surprisingly long-lasting effects on blood and tissue immunophenotypes but, importantly, bsAb treatment did not result in acute inflammatory cytokine production or persistent lymphopenia.

**Figure 2.**
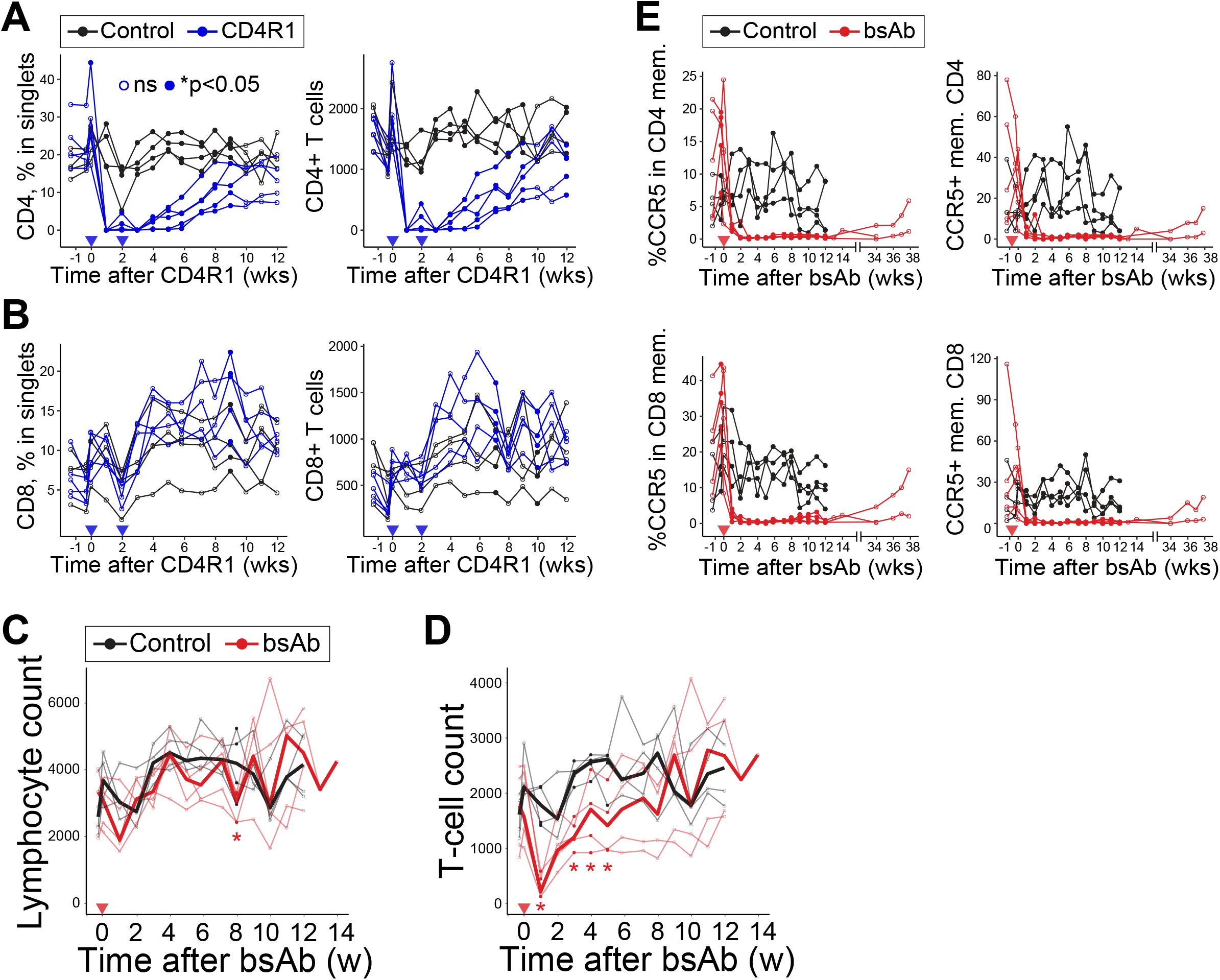
Effects of CD4R1 or CD3/CCR5 bsAb on lymphocyte subsets. **(A)** In control animals and CD4R1 recipients, CD3^+^CD4^+^ T cells detected by flow cytometry, as a percentage of singlet lymphocytes (left) or absolute counts per ml of blood (right). Linear regression was performed at each time point using the percentage values as the dependent variable and group membership as the independent variable; plotted markers are filled for each time point at which the group-membership coefficient’s p value was <0.05. The same procedure was used for all graphs in panels A, B, and E. Data for 14 animals are shown in this figure and the remaining data relevant to depletion, collected on a different schedule, in Extended Data Fig. 4. **(B)** CD3^+^CD8^+^ T cells detected by flow cytometry, as a percentage of singlet lymphocytes or absolute counts. **(C)** In CCR5/CD3 bsAb recipients, lymphocyte counts are not changed by bsAb treatment. **(D)** CD3^+^ T-cell counts. **(E)** CCR5 expression as a percentage of CD3^+^CD4^+^CD95^+^ memory T cells (upper left); absolute counts of CCR5^+^ memory CD4^+^ T cells (upper right). CCR5 expression as a percentage of CD3^+^CD8^+^CD95^+^ memory T cells (lower left); absolute counts of CCR5^+^ memory CD8^+^ T cells (lower right).

### Viral rebound delayed or prevented in some depleted macaques

ART was withdrawn after 18 weeks of therapy to assess viral rebound, defined as reappearance of >29 copies of SIV per ml of plasma in an animal that had exhibited complete suppression at the time of ART interruption. Peripheral viremia was observed after one or two weeks in all controls (Fig. 3A) and 7/8 CD4R1-treated animals (Fig. 3B), but only 3/7 bsAb-treated animals (Fig. 3C). Furthermore, there was a two-week delay in detection of viremia in two of these three bsAb-treated animals—a delay not observed in groups A or B. Therefore, following ART interruption, 1/8 CD4R1-treated animals and 4/7 bsAb-treated animals were initially aviremic. After 97 and 173 days, two bsAb recipients rebounded, leaving two animals in this group still aviremic at approximately six months after ART withdrawal. Cox regression showed that the delay in rebound of bsAb-recipient animals was significant (p=0.008; Fig. 3D) and estimated an 89% reduction in rebound hazard. Across treatment groups, seven of thirteen macaques rebounding within two weeks of ART cessation expressed Mamu-A*01, - B*08, or -B*17 (54%) while only two of five later-rebounding macaques did so (40%). There was no apparent difference between groups A, B, and C in the slope of increase in viremia when rebound did occur (p=0.56; Fig. 3E).

**Figure 3.**
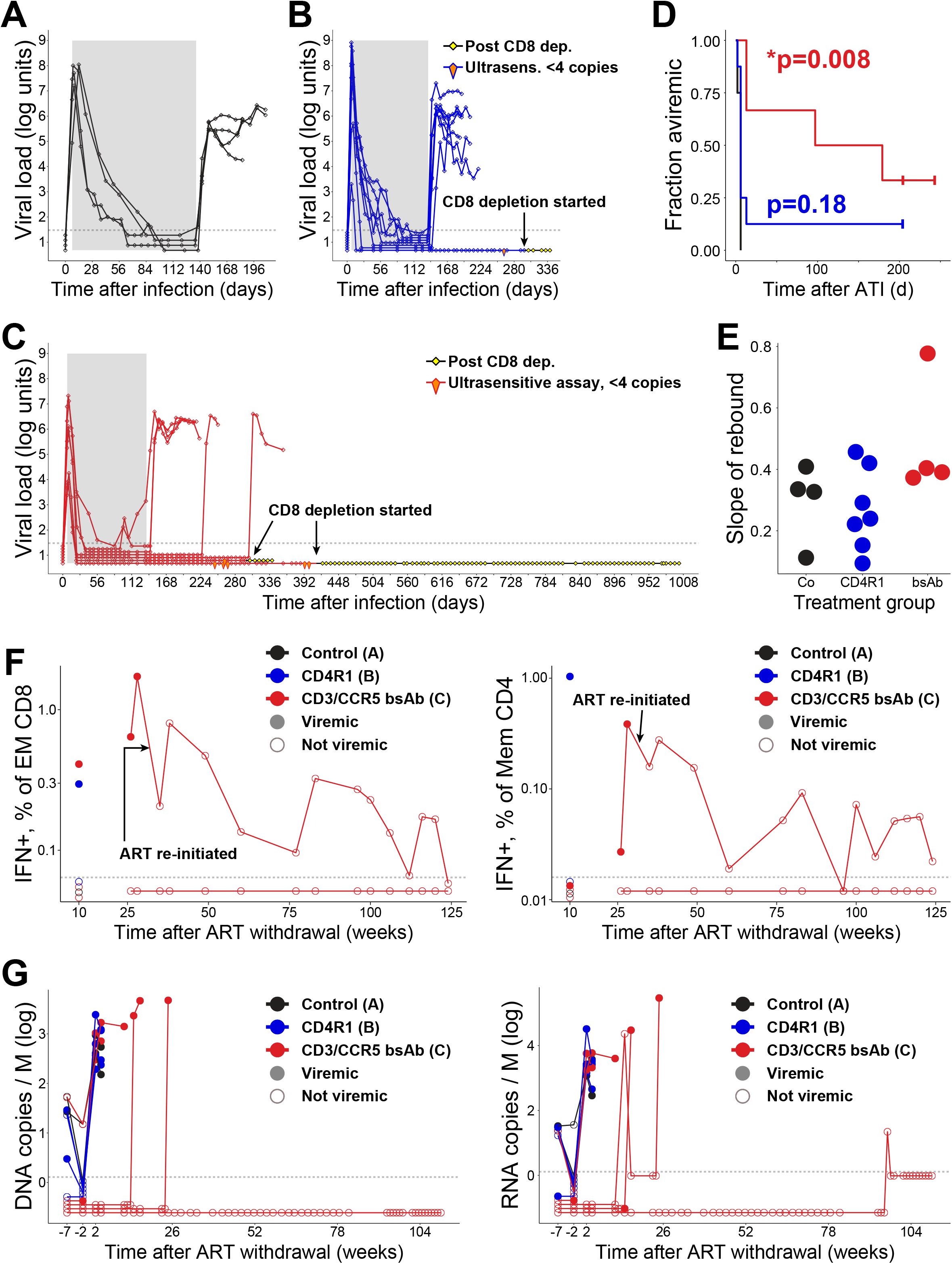
Virologic outcomes of depletion after treatment interruption. **(A-C)** Gray shading indicates the period of ART treatment. The dotted line represents the limit of detection of our viral-load assay (30 copies/ml or lower); elongated kite shapes show occasions on which high-sensitivity assays were performed, for which the limit of detection was 4 copies/ml. **(D)** Survival curves show the fraction of fully suppressed macaques in each group that maintained aviremia versus time after treatment interruption. **(E)** The slope of viremia rebound in each animal was calculated by linear regression of increasing viral loads (i.e., before the first decline, including the peak value) occurring after treatment interruption. **(F)** Immune-response assays were periodically performed using the limited cells available from these newborn macaques. Shown are IFN-γ production by CD3^+^CD8^+^CD95^+^CCR7^-^ PBMC (left) or CD3^+^CD4^+^CD95^+^ PBMC (right) after stimulation with overlapping 15-mer peptides from the SIV gag gene, assessed by cytokine flow cytometry. The figure shows results from one control macaque, two group-B (CD4R1) macaques, and five group-C (bsAb) macaques. Two of the latter animals (bsAb recipients) were tested multiple times during extended follow up. Filled markers indicate that the animal had detectable viremia at the nearest tested time point; open markers indicate that the animal did not have detectable viremia. **(G)** Cell-associated viral DNA (left) or RNA (right) copies per million PBMC. Filled markers indicate that the animal had detectable viremia at the nearest tested time point; open markers indicate that the animal did not have detectable viremia. All samples plotted below the dotted line had no detectable nucleic acid.

We next wished to determine the extent of viral clearance, as defined by absence of detectable viremia and failure to detect replication-competent virus in any sample, among animals with durable post-ART control (>6 months). First, the single CD4R1-treated and two bsAb-treated animals in remission were tested for residual viremia. No virus was detected using a high-sensitivity qPCR assay having a limit of detection of four copies/ml (orange kites in Figs. 3B-C). Second, to provoke outgrowth of any replication-competent virus residing in RCVRs, CTLs in all three animals were depleted using the rhesusized anti-CD8ɑ monoclonal antibody, MT807R1. Post-treatment FACS analysis demonstrated complete depletion of blood CD8^+^ T cells in all animals (Extended Data Fig. 3). Despite this transient hobbling of the CTL response, rebound did not occur, suggesting that CD8^+^ T-cells were not the primary mechanism of control (black/yellow diamonds in Figs. 3B-C). Indeed, T-cell responses to SIV were associated with plasma viremia and generally not measurable in controlling animals (Fig. 3F). Finally, we evaluated cell-associated SIV nucleic acid in PBMC before and after treatment interruption in a subset of animals for which sufficient cells were available, including two controls, four CD4R1 recipients, and five bsAb recipients. As expected, cell-associated DNA was seen to decline between weeks 12 and 17 after infection (weeks -7 and -2 relative to ART stop), presumably reflecting continued effective suppression by ART (Fig. 3G). Withdrawal of ART, in fact, led to an almost immediate reappearance of or dramatic increase in cell-associated SIV DNA in 8 of 11 animals, exclusive of three bsAb recipients (Fig. 3G, left, measurements two weeks post ART withdrawal). SIV DNA was first detected in two of these latter animals at 16 or 25 weeks post ART withdrawal. In Fig. 3G, filled symbols represent detectable viremia, so it is apparent that the animals with detectable SIV DNA at 16 or 25 weeks are the late rebounders in Fig. 3C. In one aviremic animal (still in remission) cell-associated SIV DNA was not measurable at any point in the follow up period. Cell-associated SIV RNA (Fig. 3G) demonstrated similar dynamics, with decline between 12 and 17 weeks after ART initiation, and 8/11 animals manifesting RNA very soon after ART withdrawal. Two bsAb recipients were delayed in RNA production and, interestingly, in one case measurable PBMC-associated viral RNA preceded detectable viremia by many weeks. Remarkably, only one instance of RNA detection (2 copies in 913,000 cells), without DNA or viremia, occurred at week 97 in the macaque that remains aviremic today, over three years following ART withdrawal. In sum, these data provide remarkable evidence that CD4 or CCR5 depletion in recently infected infants can profoundly alter both the kinetics of SIV spread, as reflected in viremia, and RCVR establishment, such that 1/8 CD4R1-treated and 2/7 bsAb-treated animals achieved remission that was maintained for >6 months.

### RCVR depletion achieved by bispecific antibody

Clinical experience and mathematical modeling have shown that indefinite control over viremia is an attractive but problematic experimental endpoint, because the RCVR can be undetectable and still cause rebound after prolonged aviremia. We chose to follow one of the three aviremic, previously CD8-depleted macaques long term (a bsAb recipient) and to necropsy the other two animals in order to rigorously search for persisting virus. At necropsy, bone marrow, intestine (duodenum and colon), lung, spleen, and multiple lymph nodes (cervical, axillary, inguinal, mesenteric) were harvested for qPCR and RNAscope. In contrast to bsAb-treated animals with rebound following ART interruption and to untreated control animals, no residual SIV RNA or DNA was detected by qPCR in any of the assayed tissues (Fig. 4A). Similarly, an RNAscope survey revealed no viral genomes in the tissues of the aviremic bsAb-treated macaque that was necropsied, while an identical survey of a viremic, bsAb-treated animal revealed unambiguous SIV staining in colon, lymph node, and spleen (Fig. 4B). Viral outgrowth assays from lymph-node cells failed to recover SIV in 8 of 8 attempts from the two non-rebounding and necropsied animals (Fig. 4C; 4.7 × 10^7^ or 3.4 × 10^7^ total cells tested), while lymph-node biopsies from two control animals did grow SIV both when the animals were viremic and when they were not.

**Figure 4.**
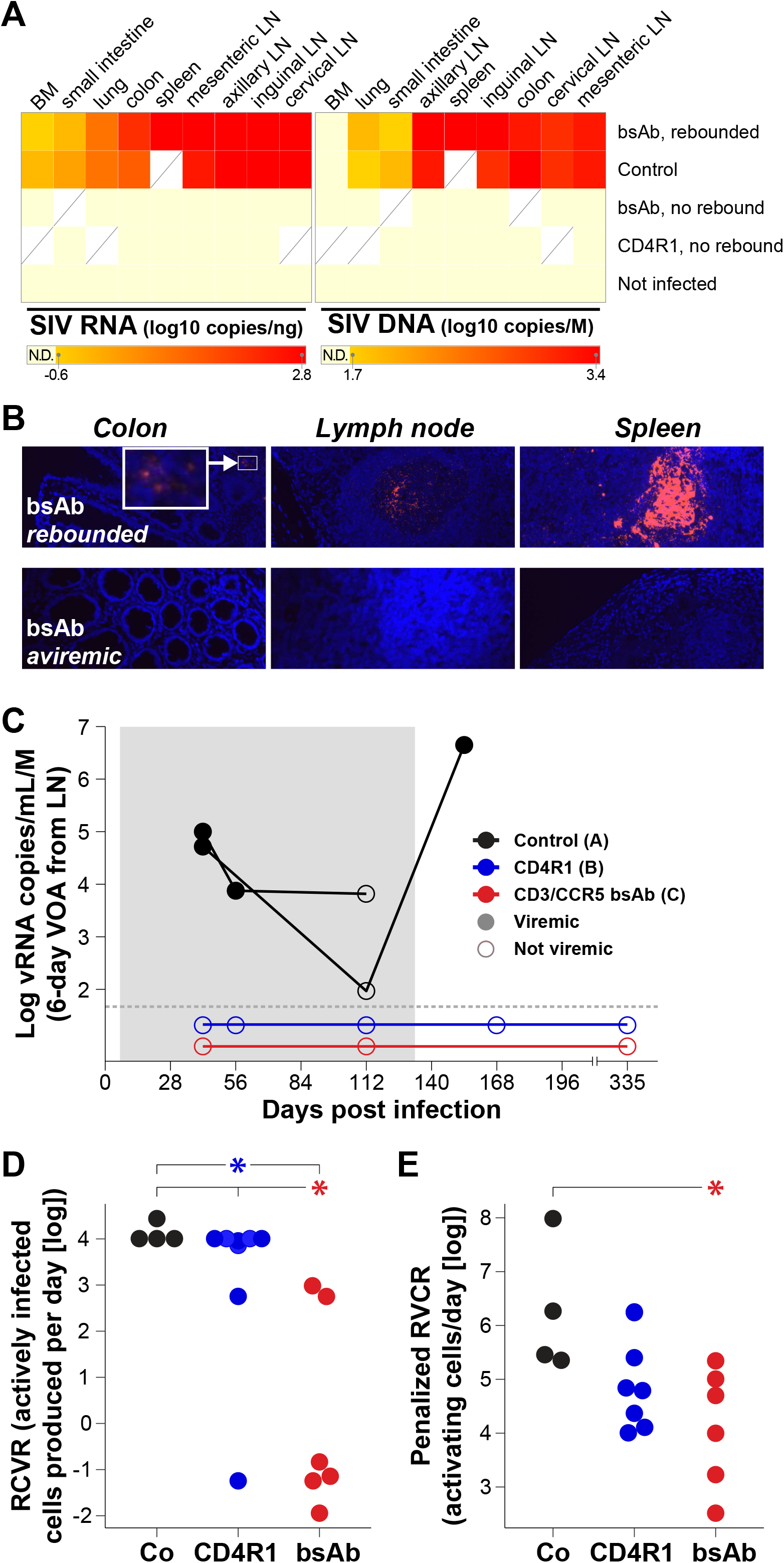
Immune responses and tests for residual virus in tissue samples. **(A)** Quantitative PCR results from the tissues of one bsAb-treated and one CD4R1-treated animal that did not rebound during the period of follow up and were necropsied (rows 2-3, “no rebound”), along with results from positive or negative controls. Viral nucleic acid was detected in all tissues of a necropsied control animal (second row) or a necropsied bsAb recipient that did rebound (first row), but not in any tissues of the non-rebounding bsAb and CD4R1 recipients. **(B)** RNAscope analysis of colon, lymph node, and spleen tissues from two animals in group B again detected viral nucleic acid only in a rebounding bsAb recipient. **(C)** Viral outgrowth was detected from lymphocytes of control macaques when the animals were viremic (filled markers) or aviremic (open markers). No outgrowth was detected from eight lymphocyte samples from depleted macaques that never rebounded. **(D)** Estimation of the size of the viral reservoir prior to interruption of ART. **(E)** Moderation of the estimate in viral reservoir size by adjusting for peak virus load.

Finally, to gain quantitative insight into the reduction in the RCVR provided by CD4R1 or CCR5/CD3 bsAb treatment, we used the survival curves for rebound viremia and the statistical tools published by Hill and colleagues.^16,17^ These tools integrate viral load and viral outgrowth data to provide information about the likely size of the underlying RCVR. Using this model to calculate most-likely RCVR sizes for each macaque (measured in terms of the rate at which the RVCR produces actively infected cells), we found that likely RCVRs in both CD4R1- and bsAb-treated macaques at the time of ART interruption were smaller than those in control animals (Fig. 4D; p=0.01 and 0.0002, respectively, by Kruskal-Wallis and Conover-Iman testing). The estimated RCVR in bsAb recipients was also significantly smaller than that in CD4R1 recipients (Fig. 4D; p=0.005). Next, we moderated these estimates of the RCVR at ART interruption to correct for the possibility of lesser RCVR establishment in macaques having lower viral loads before treatment initiation. Briefly, calculated RCVRs at ART interruption were inflated for all animals having lower viremia peaks, by the corresponding number of log units, to account for the possibility that calculated reservoir sizes were low due to reduced early viral replication. In this penalized analysis, the median reservoir in bsAb-treated macaques remained significantly smaller than that in control animals (Fig. 4E; p=0.005) but not in CD4R1 recipients (p=0.1).

## Discussion

We found that newborn macaques receiving depletion therapy can achieve durable remission, including aviremia that is maintained for up to 3.25 years following treatment interruption and despite depletion of CD8^+^ T cells. This hopeful finding is surprising for many reasons. Mathematical modeling suggests that a 2,000-fold reduction in the reservoir is required to delay rebound by one year in the majority of treated people,^16,17^ while a 5,000-fold reduction is required for a 50% chance of sustaining remission for three years. An efficiency of depletion of even 99.9% in peripheral blood and less in tissues^14^ would seem unequal to the task. One possibility is that CCR5 depletion alters key parameters that control viral rebound, such as the rate at which virus from reactivated cells can infect of other targets, or even the likelihood of reactivation from latency. Complete elimination of all latently infected cells is perhaps unnecessary because under some circumstances it appears host defenses can eliminate nascent infectious foci,^18^ and host immunity may also contribute to restriction of most HIV transmission events to single transmitted founders.^19^ With respect to transmission, it seems likely that many host cells are infected with replication-competent virus but that the vast majority are eliminated by innate or adaptive immune responses before dissemination. Innate-immune sieving has been amply demonstrated,^20^ while adaptive-immune sieving would be consistent with the resistance of animals inheriting “controller” MHC to oral challenge that is described above.

Apparent clearance of the RCVR via CCR5 targeting alone is particularly surprising because, although CCR5 is needed for infection by most SIV variants, it is not a known marker of the HIV reservoir. The reservoir under ART is concentrated among central memory (T_CM_) and transitional memory (T_TM_) CD4^+^ T cells.^21^ More recently, a less differentiated subset of long-lived cells with high self-renewal capacity, the stem-cell memory CD4^+^ T cell (T_SCM_), has been identified as a contributor to long-term HIV persistence.^22,23^ The persistent reservoir has also been described according to T-cell function and homing capacity: Th17 and Th1/Th17 CD4^+^ T cells, as well as cells expressing CCR6 and CXCR3, show increasing contribution to the reservoir with duration of ART.^24,25^ Chomont and colleagues found that CD4^+^ T cells expressing PD-1, TIGIT and LAG-3 alone or in combination are enriched for persistent HIV during ART.^26^ Others reported that CD32a is a marker of the CD4 T-cell HIV reservoir harboring replication-competent proviruses,^27^ but this finding has been debated.^28^

Notably, however, this field-wide examination of the HIV reservoir has been focused on chronically infected individuals. CCR5 may be a good marker of the RCVR in early infection because CCR5^+^ cells are the first to be infected^29–31^ and because some of these CCR5^+^ cells, including CCR5^+^ pre-Tfh, can cease expression in the process of differentiation to longer-lived cells that contribute to the RCVR.^32^ Another possibility is that CCR5 depletion acts not only through elimination of infected cells, but also through depletion of cells that are vulnerable to infection by re-activated SIV, i.e., new targets. In this conception, remaining cells in the RCVR may occasionally re-activate and produce virus, but this virus finds no foothold among remaining immune cells, while the RCVR is gradually diminished due to toxicity of viral replication and/or attrition of the cells with time. It is additionally possible that bsAb serves as an entry inhibitor by preventing access of the virus to its co-receptor. If so, then blockade of CCR5 with binding antibodies and/or small molecules may raise the chance of post-treatment cure.

Possibly, such drugs could even potentiate the bsAb effects demonstrated here, leading to more frequent cases of remission than we observed in this study. The extent and durability of CCR5 depletion achieved in this study with a single administration of bsAb were unexpected. CCR5 expression has been associated with T-cell activation,^33–36^ suggesting that CCR5 expression is dynamic and therefore unlikely to be susceptible to long-term depletion with a single dose of drug. An early detailed study of CXCR4 and CCR5 regulation in response to activation of human T cells, however, showed that CCR5 expression was unchanged throughout 12 days of continuous stimulation with PHA or anti-CD3.^37^ CCR5 expression was found to be up-regulated after nine days of IL-2 treatment. A recent detailed review described unexpectedly broad CCR5 expression across many human CD4^+^ T-cell subsets including Th1, Th17, and T-reg cells and on CD8^+^ T cells expressing Granzyme K.^38^ In both humans and macaques, some PD-1^high^CXCR5^+^ Tfh express CCR5, which could be key to the success of anti-CD3/CCR5 in RCVR depletion.^38,39^ It is clear that presence of CCR5 on the cell surface can be reduced by receptor internalization due to ligand binding.^40^ Nonetheless the durability of the depletion we observed suggests that CCR5 expression is primarily a stable characteristic of certain cell lineages.

Adults with the CCR5 Δ32/Δ32 homozygous genotype have not shown any obvious immunodeficiency, apart from reports of increased susceptibility to symptomatic West Nile virus encephalitis.^41–43^ This apparent tolerance of lifelong CCR5 deficiency suggests that transient depletion of CCR5-expressing cells to achieve HIV cure may be an acceptable risk. Our study demonstrates that targeting CCR5-expressing cells using the host immune system may be a surprisingly effective route to HIV cure, at least for individuals that can be treated soon after infection. Furthermore, the pronounced delay in rebound that we observed following depletion with CD3/CCR5 bsAb might reinforce or potentiate other approaches to cure.

## TABLES

None

## METHODS

### Experimental animals and samples

This study was approved in advance by the University of California, Davis (UC Davis) Institutional Animal Care and Use Committee (IACUC) and was performed at the California National Primate Research Center (CNPRC). Housing, medical care, and all procedures were performed in accordance with UC Davis IACUC-established policies. UC Davis has an Animal Welfare Assurance on file with the NIH Office of Laboratory Animal Welfare and is fully accredited by the Association for the Assessment and Accreditation of Laboratory Animal Care International. Twenty-four newborn rhesus macaques (*Macaca mulatta*) were selected from the conventional colony at CNPRC. All animals tested negative for HIV-2, SIV, simian T cell lymphotropic virus type 1 (STLV), and type-D retrovirus at the start of the study. Animals were housed at the CNPRC. Animals were administered 10 mg/kg body weight ketamine-HCl intramuscularly when necessary for immobilization. Analgesics were administered at the discretion of the CNPRC veterinary staff to minimize pain and discomfort. Plasma and peripheral blood mononuclear cells (PBMC) were prepared from EDTA anticoagulated blood and stored at -80°C as previously described.^1^ Mononuclear cells from lymph node biopsies were also prepared and stored as previously described.^1^

### Virus and cART

Macaques were challenged orally with 50,000 TCID_50_ of SIVmac251 twice on the day of inoculation as previously described.^2^ Challenge virus was provided by the CNPRC Virology Core. Antiretroviral triple therapy, consisting of tenofovir disoproxil fumarate (5.1 mg/kg), emtricitabine (40 mg/kg), and dolutegravir (2.5 mg/kg), was delivered subcutaneously each day.

### Cytotoxic antibodies

Anti-CD3/CCR5 bispecific antibody, produced and purified as previously described,^3^ was infused intravenously at 3 mg/kg. Rhesus IgG1 recombinant anti-CD4 [CD4R1] antibody was acquired from the NIH nonhuman primate reagent resource (NHPRR) and administered at 50 mg/kg by intravenous infusion according to the manufacturer’s instructions. Animals were pretreated with diphenhydramine to reduce the risk of anaphylactoid reactions to the antibody therapy. Oxygen was administered post-infusion at the veterinarian’s discretion.

### Immunophenotyping and whole blood staining

PBMC and lymph node cells were stained and analyzed by flow cytometry as previously described.^3^ Absolute cell subset counts were obtained with dual-platform techniques utilizing flow cytometry to provide the percentage of a given cell subset among total blood leukocytes along with a hematology analyzer. EDTA-anticoagulated whole blood was assessed with a ABX Pentra 60 C+ hematology analyzer (HORIBA) or stained with monoclonal antibodies against cell markers including CD3 (clone SP34-2), CD4 (clone L200), CD8 (clone SK1), CD11c (clone N418), CD11b (clone ICRF44), CD14 (clone M5E2), CD16 (clone 3G8), CD20 (clone 2H7), CD25 (clone M-A251), CD28 (clone CD28.2), CD80 (L307.4), CD95 (clone DX2), CD159 (NKG2A, clone REA110), CD185 (CXCR5, clone SPRCL5), CD195 (CCR5, clone 3A9), CD279 (PD-1, clone EH12.2H7), and HLA-DR (clone G46-6). The anti-CCR5 clone, 3A9, binds to a different epitope than the clone used for bsAb construction (5C7); the two antibodies were shown to bind simultaneously to CCR5-expressing cells. All fluorochrome-conjugated monoclonal antibodies were purchased from BD Biosciences, BioLegend, Thermo Fisher Scientific, or Miltenyi Biotec. Lysis and fixation of whole blood samples were performed by the TQ-Prep Workstation and IMMUNOPREP reagent system (Beckman Coulter). Cells were acquired on a BD FACSymphony A3 Cell Analyzer (BD Biosciences) and data were analyzed using FlowJo software (BD Biosciences).

### Cytokine flow cytometry assay

Levels of SIV-specific cytokine-producing cells were determined by flow cytometry after stimulation with sequential 15-mer peptides (overlapping by 11 amino acids) comprising the SIVmac239 Gag, Rev, or Nef (provided by NIH AIDS Reagent Program) and anti-CD28 (clone CD28.2, NIH Nonhuman Primate Reagent Resource) and anti-CD49d (clone L25, BD Biosciences) monoclonal antibodies. Cells were incubated with peptides (or DMSO control) and co-stimulatory antibodies for 1 h, followed by adding Brefeldin A (Sigma-Aldrich) for an additional 8 h. Cells were stained for surface markers with LIVE/DEAD Fixable Aqua Dead Cell Stain Kit (Thermo Fisher Scientific), anti-CD3-APC-Cy7, anti-CD4-BV650, anti-CD8-PerCP-Cy5.5, anti-CD95-APC, anti-CCR7-BV421 (clone 3D12). Cells were then washed, permeabilized using a Cytofix/Cytoperm solution kit (BD Biosciences), and intracellularly stained with anti-TNF-Alexa 700 (alone MAB11), and anti-IFN-γ-PE-Cy7 (clone B27). After staining, cells were washed, fixed with 1% paraformaldehyde in staining buffer (PBS containing 1% FBS), and acquired using a LSRFortessa cell sorter operated by FACSDiva software (BD Biosciences).

### CD8 T cell depletion

Rhesusized anti-CD8 alpha antibody [MT807R1]^4,5^ acquired from the NHPRR was used to deplete CD8+ lymphocytes. The antibody was administered according to manufacturer’s instructions (SOP A06-01); at 10 mg/kg subcutaneously on day 0, and at 5 mg/kg intravenously on days 3, 7, and 10 of the treatment protocol.

### Viral detection assays

SIV RNA in plasma was measured by quantitative RT-PCR as described.^6,7^ Key tissues collected at necropsy were assessed for viral RNA and DNA. Tissue samples were stored in RNA Later (Invitrogen) at -20°C until purification and quantitative PCR analysis. RNA and DNA were purified using a Qiagen All Prep kit according to manufacturer’s instructions and stored at -20°C. For RNA, cDNA was generated as previously described.^8^ The cDNA and purified viral DNA were used for quantitative PCR (qPCR) using QuantiTect probe PCR master mix (Qiagen) and optimized concentrations of forward primer (TGTGTGGGAGACCATCAAGC), reverse primer (GCTCCCTAAGTTGTCCTTGTTG’), and probe (FAM/ACGAGGAGGCTGCAGATTGGGACTTGCA/NFQ/MGB-3’) in a QuantStudio 12K Flex real-time cycler (Applied Biosystems). PCR amplicons of the SIVgag target were used to generate a standard curve and line equation for qPCR analysis. PBMC collected longitudinally were also assessed for viral RNA and DNA. Cells were lysed with one milliliter TRIzol Reagent (Ambion) and RNA and DNA were purified sequentially from half of the lysate using a Direct-zol DNA/RNA Miniprep kit (Zymo) according to the manufacturer’s protocol. Cell-associated SIV RNA was quantified using a QuantStudio 6 Flex instrument (Applied Biosystems) and a Superscript III Platinum One-Step qRT-PCR System (ThermoFisher). Primers and reporter probe bind to the LTR and their sequences are GCTAGACTCTCACCAGCACTTG (forward), CTAGGAGAGATGGGAACACACA (reverse) and 56-FAM/TCCACGCTT/ZEN/GCTTGCTTAAAGCCCTC/3IABkFQ (probe). Cycling consisted of 50°C for 15 m, 95°C for 2 m followed by 45 repeats of 95°C for 15 s and 62°C for 30 s. To measure SIV DNA, the same reagents were used with Taqman Fast Universal PCR Mastermix (Applied Biosystems) and the same cycling except for the 50°C cDNA synthesis step. The single-copy C-C chemokine receptor type 5 (CCR5) gene was quantified to determine the number of cell equivalents in DNA samples for standardization purposes. Reagents to detect CCR5 are GTCTTCATTACACCTGCAGCTCTCA (forward), AGGATTCCCGAGTAGCAGATGA (reverse) and 56-FAM/AGCAGCGGC/ZEN/AGGACCAGCCCCAAG/3IABkFQ (probe) and cycling was the same as that used to amplify SIV DNA.

### SIV RNA detection in tissues by ISH

Tissues collected at necropsy were analyzed for SIV RNA by *in situ* hybridization (ISH) using an Advanced Cell Diagnostic (ACD) RNAScope Fluorescent Multiplex Kit V2 for rhesus macaques according to the manufacturer’s instructions. Tissues fixed in 10% neutral buffered formalin were embedded in paraffin and cut into 5 μm sections. SIV RNA was detected using a previously described custom probe.^9^ Probes supplied with the RNAScope kit (ACD) specific for peptidylprolyl isomerase B (PPIB) and *dapB* sequences were used as positive and negative controls, respectively. Following Tyramide Signal Amplification (TSA) using a cyanine-5 kit (Cy5; Perkin Elmer) and DAPI staining, slides were mounted with ProLong Gold (ThermoFisher). Slides were cured and then images were captured on an Evos2 Auto Fluorescent System. Images were analyzed using FIJI (Fiji is Just ImageJ) and its macro language for automated segmentation, deconvolution, and quantification.^10^

### Viral outgrowth assays

Mononuclear lymph node cells were mixed with 174xCEM cells and activated for 24 hours in RPMI 1640 growth media containing fetal bovine serum (FBS; 10%), interleukin-2 (IL-2; 50 U/mL), phytohaemogglutinin (PHA; 5 μg/mL), and ionomycin (1 μg/mL). After 24 hours, PHA and ionomycin were removed and cultures were maintained for 28 days or until cytopathic effect was apparent in 174xCEM cells. Cell-free cell supernatants were collected and stored at -80°C. Viral RNA was purified from the supernatants using a Qiagen UltraSens virus kit according to the manufacturer’s instructions. cDNA was generated and used for qPCR as described above.

### Estimation of probable RCVR at ART interruption

We applied the formulas and modeled survival curves from refs. 11 and 12 to derive posterior probabilities for the “latent reservoir exit rate” when treatment is stopped, i.e., the number of actively infected cells produced per day by the remaining RCVR (expressed in the variable, *A*). Briefly, for macaques that do not rebound to a given time, the probability distribution is given as a normalized product of (i) the likelihood of survival to that time, given possible values of *A* (from the survival curves in ref. 12 and [http://www.danielrosenbloom.com/reboundtimes]), and (ii) the prior probability distribution for those values of *A* (uniform or given by results of viral-outgrowth assays as in ref. 13). For macaques that did rebound, the formula is slightly modified so that the likelihood function for the time of rebound is taken from the slopes of the survival curves at that time. As in ref. 11, we then determine a point estimate for each animal by taking the median of the posterior distribution (for macaques that rebounded) or the largest value of *A* for which the likelihood of survival to the follow-up time exceeds 0.5 (for those not rebounding).

## END NOTES

## Acknowledgements

We would like to thank Anthony Michaels, Rijian Wang, Carolyn Kraus, and Joanna Zikos for production and purification of the CCR5/CD3 bi-specific antibody. We would also like to thank the veterinarians and staff at the CNPRC. This work was supported with funding from the California HIV/AIDS Research Program (Award No. ID13-D-564 [DJHO]); National Institute of Allergy and Infectious Diseases, National Institutes of Health (NIH) (Award Nos. R21 AI116230, R01 AI143554, and R01 AI150554 [DJHO]); and National Cancer Institute, NIH (Contract No. 75N91019D00024/HHSN261201500003I [JDL]).

## Author Contributions

JDD, DM, KML, and DJHO designed and managed the study. GML and KML assessed cellular depletion. KML, WLWC, and HK performed functional T cell assays. WLWC and HK performed Luminex assays. JS and MS performed qPCR for RNA and DNA from cell pellets post-ART withdrawal. JD and JF performed qPCR on tissue samples. DM performed RNAscope. JDD performed viral outgrowth assays. DJHO performed mathematical modeling. CNM provided critical feedback and support in data interpretation. KE, DM, and KR designed and generated the therapeutic antibodies. WL prepared and managed combination antiretroviral therapy. JDD and HK prepared inoculations and antibody injections. HK, JF, and WL processed blood samples. JDD, HK, JD, JF, WL, NC, and CC assisted with necropsies. JDL oversaw plasma SIV RNA measurements performed in the Quantitative Molecular Diagnostics Core of the AIDS and Cancer Virus Program of the Frederick National Laboratory. JDD and DJHO wrote the manuscript, with all authors providing editorial support.

## ADDITIONAL INFORMATION

## EXTENDED DATA FIGURE/TABLE LEGENDS

**Extended Data Figure 1.**
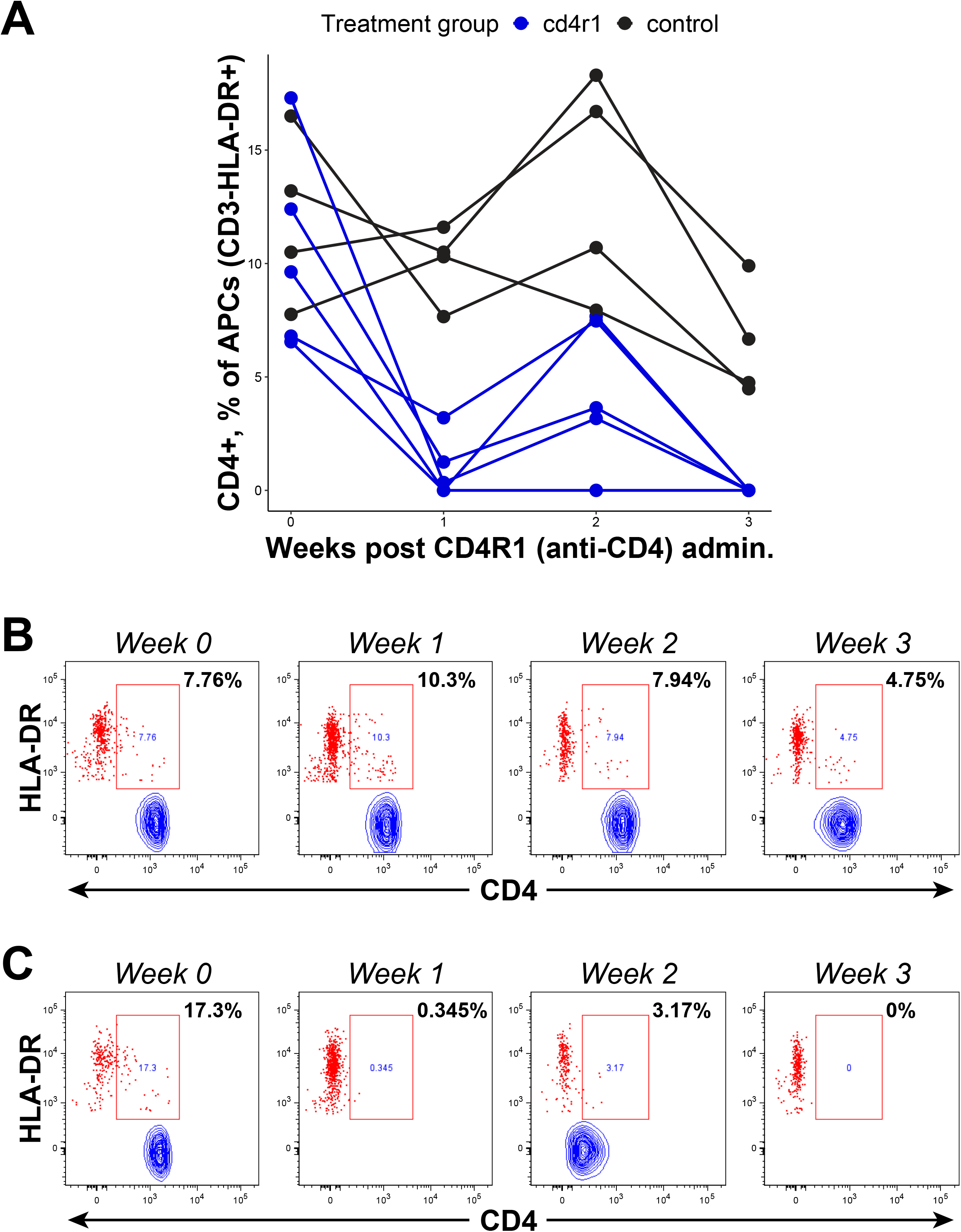
Depletion of CD4-expressing APCs among CD4R1-treated macaques. **(A)** After a first administration of depleting antibody (week 0), representation of CD4-expressing cells among APCs declines precipitously. After a second administration (week 2), depletion of CD4^+^ APCs is complete. **(B)** Maintenance of CD4 expression by the APCs of a control (undepleted) animal. **(C)** Complete depletion of CD4-expressing APCs by the fourth week after anti-CD4 antibody administration.

**Extended Data Figure 2.**
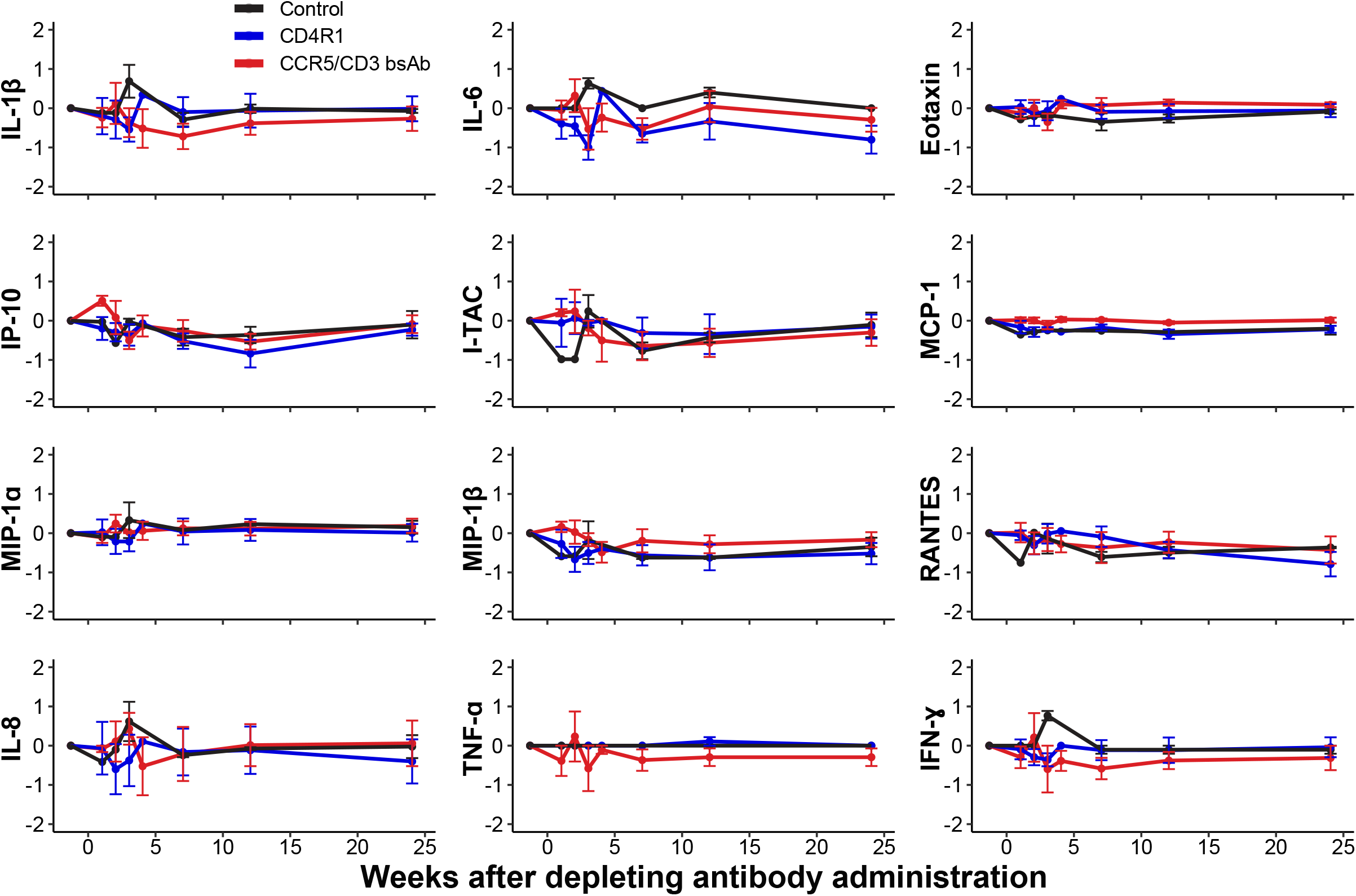
No significant effect of CD4 or CCR5 depletion on cytokines in blood. Change in the log concentrations of 12 cytokines measured by Luminex are shown. Group averages are plotted with error bars indicating the standard error. There are no significant differences in cytokine concentration that are attributable to depletion using either CD4R1 (administered days 0 and 14 on the x axis) or CD3/CCR5 bsAb (day 0 only).

**Extended Data Figure 3.**
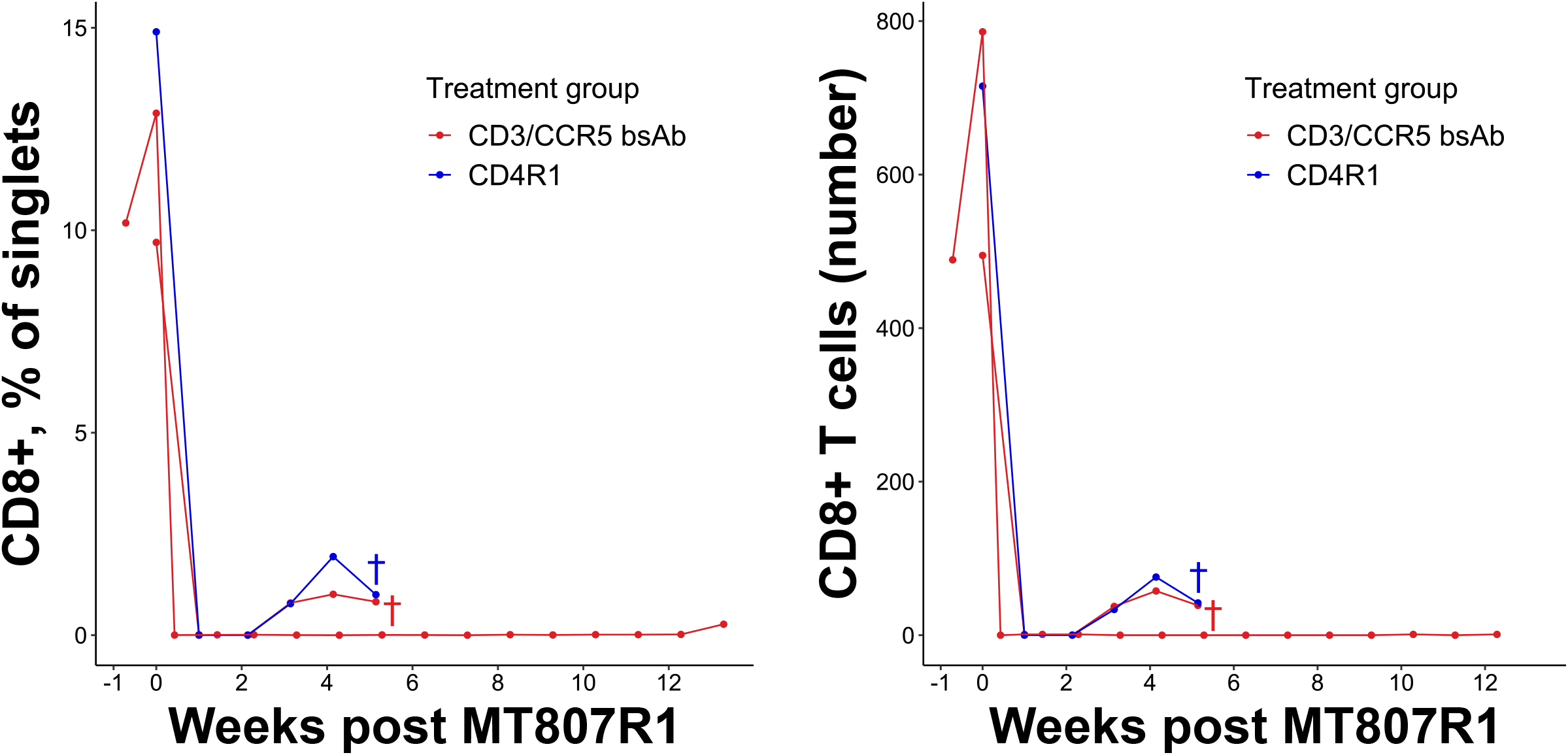
Depletion of CD8+ T cells from macaques treated with CD4R1 or CD3/CCR5 bsAb that were exhibiting post-treatment control. One CD4R1 recipient and two bsAb recipients were treated. The rhesus IgG1 recombinant anti-CD8 alpha (MT807R1) monoclonal antibody was engineered and produced by the Nonhuman Primate Reagent Resource (NIH Nonhuman Primate Reagent Resource [NHPRR] Cat. #PR-0817, RRID:AB_2716320). The antibody was administered in four doses on days 0, 3, 7, and 10 according to NHPRR instructions. The crosses show when two animals were necropsied to allow testing of tissue for residual virus (Fig. 4).

**Extended Data Figure 4.**
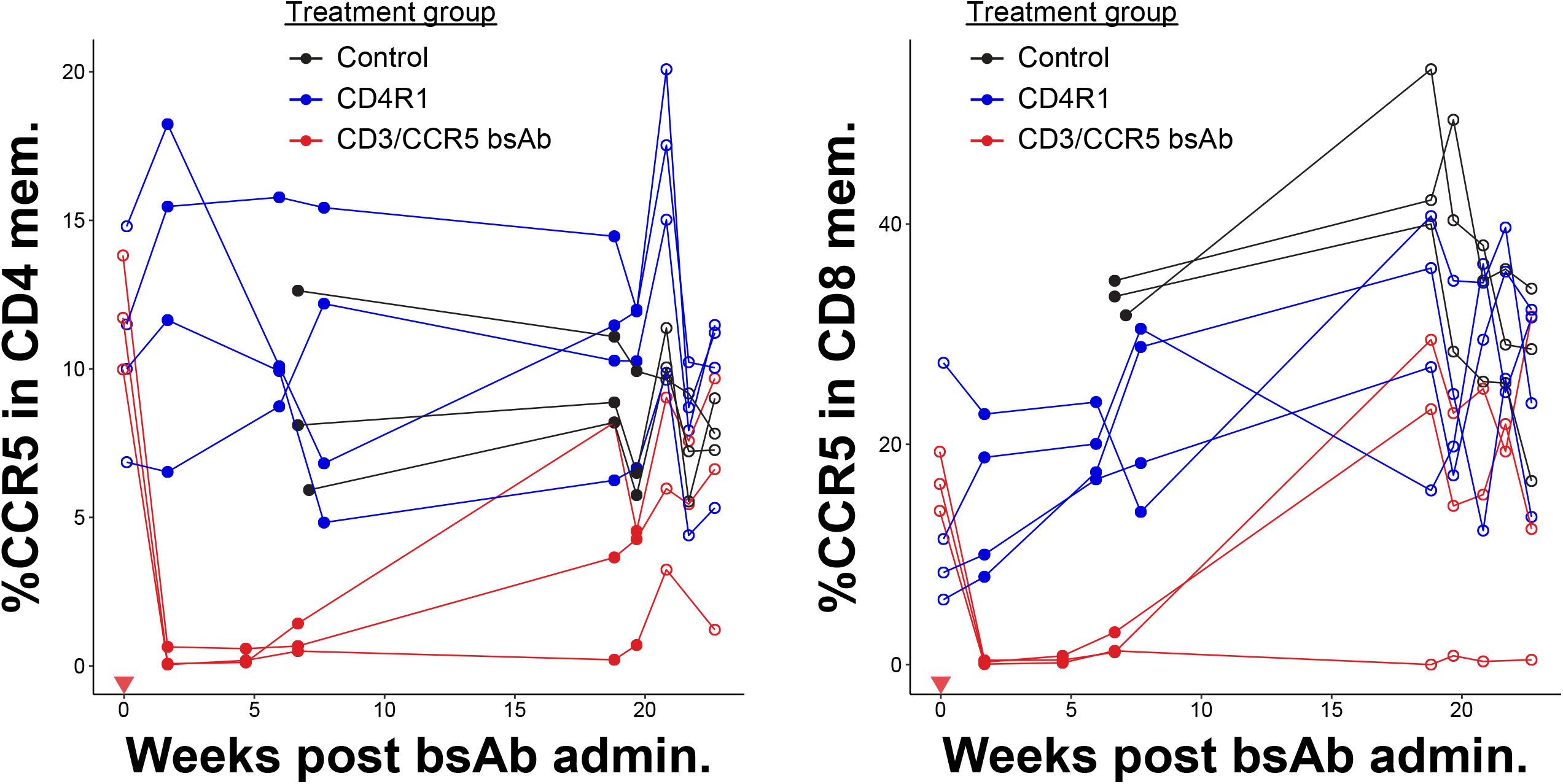
Effects of CD3/CCR5 bsAb on CCR5-expressing cells in the CD4 and CD8 compartments. CCR5 expression as a percentage of CD3^+^CD4^+^CD95^+^ memory T cells (left) or as a percentage of CD3^+^CD8^+^CD95^+^ memory T cells (right). Filled markers indicate a significant difference in the percentage of CCR5^+^ cells at that time point in bsAb-recipient vs. control or CD4R1-recipient macaques.

## MAIN REFERENCES

1. Hütter, G. et al. Long-Term Control of HIV by CCR5 Delta32/Delta32 Stem-Cell Transplantation. New England Journal of Medicine 360, 692–698 (2009).

2. Persaud, D. et al. Absence of Detectable HIV-1 Viremia after Treatment Cessation in an Infant. New England Journal of Medicine 369, 1828–1835 (2013).

3. Bertolli, J. et al. Estimating the Timing of Mother-to-Child Transmission of Human Immunodeficiency Virus in a Breast-Feeding Population in Kinshasa, Zaire. J Infect Dis 174, 722–726 (1996).

4. Menu, E. et al. Selection of maternal human immunodeficiency virus type 1 variants in human placenta. European Network for In Utero Transmission of HIV-1. J Infect Dis 179, 44–51 (1999).

5. Emau, P. et al. Post-exposure prophylaxis for SIV revisited: Animal model for HIV prevention. AIDS Res Ther 3, 29 (2006).

6. Kahn, J. O. et al. Feasibility of postexposure prohpylaxis (PEP) against human immunodeficiency virus infection after sexual or injection drug use exposure: the San Francisco PEP Study. J Infect Dis 183, 707–714 (2001).

7. Roland, M. E. et al. Seroconversion Following Nonoccupational Postexposure Prophylaxis against HIV. Clinical Infectious Diseases 41, 1507–13 (2005).

8. Whitney, J. B. et al. Rapid seeding of the viral reservoir prior to SIV viraemia in rhesus monkeys. Nature 512, 74–77 (2014).

9. Borducchi, E. N. et al. Antibody and TLR7 agonist delay viral rebound in SHIV-infected monkeys. Nature 563, 360–364 (2018).

10. Okoye, A. A. et al. Early antiretroviral therapy limits SIV reservoir establishment to delay or prevent post-treatment viral rebound. Nat Med 24, 1430–1440 (2018).

11. Ndung’u, T., McCune, J. M. & Deeks, S. G. Why and where an HIV cure is needed and how it might be achieved. Nature 576, 397–405 (2019).

12. Li, C., Mori, L. & Valente, S. T. The Block-and-Lock Strategy for Human Immunodeficiency Virus Cure: Lessons Learned from Didehydro–Cortistatin A. J Infect Dis 223, S46–S53 (2021).

13. Saag, M. S. et al. Antiretroviral Drugs for Treatment and Prevention of HIV Infection in Adults: 2020 Recommendations of the International Antiviral Society– USA Panel. JAMA 324, 1651–1669 (2020).

14. Merriam, D. et al. Depletion of Gut-Resident CCR5+ Cells for HIV Cure Strategies. AIDS Res Hum Retroviruses 33, S70–S80 (2017).

15. Iacob, S. A. & Iacob, D. G. Ibalizumab Targeting CD4 Receptors, An Emerging Molecule in HIV Therapy. Front Microbiol 8, 2323 (2017).

16. Hill, A. L., Rosenbloom, D. I. S., Fu, F., Nowak, M. A. & Siliciano, R. F. Predicting the outcomes of treatment to eradicate the latent reservoir for HIV-1. Proc Natl Acad Sci U S A 111, 13475–13480 (2014).

17. Hill, A. L. et al. Real-Time Predictions of Reservoir Size and Rebound Time during Antiretroviral Therapy Interruption Trials for HIV. PLoS Pathog 12, e1005535 (2016).

18. Hansen, S. G. et al. Immune clearance of highly pathogenic SIV infection. Nature 502, 100–104 (2013).

19. Keele, B. F. et al. Identification and characterization of transmitted and early founder virus envelopes in primary HIV-1 infection. Proc Natl Acad Sci U S A 105, 7552–7557 (2008).

20. Gondim, M. V. P. et al. Heightened resistance to host type 1 interferons characterizes HIV-1 at transmission and after antiretroviral therapy interruption. Sci. Transl. Med 13, 8179 (2021).

21. Chomont, N. et al. HIV reservoir size and persistence are driven by T cell survival and homeostatic proliferation. Nat Med 15, 893–900 (2009).

22. Buzon, M. J. et al. HIV-1 persistence in CD4+ T cells with stem cell–like properties. Nat Med 20, 139–142 (2014).

23. Jaafoura, S. et al. Progressive contraction of the latent HIV reservoir around a core of less-differentiated CD4+ memory T Cells. Nat Commun 5, 1–8 (2014).

24. Khoury, G. et al. Persistence of integrated HIV DNA in CXCR3 + CCR6 + memory CD4R T cells in HIV-infected individuals on antiretroviral therapy. AIDS 30, 1511–1520 (2016).

25. Sun, H. et al. Th1/17 Polarization of CD4 T Cells Supports HIV-1 Persistence during Antiretroviral Therapy. J Virol 89, 11284–11293 (2015).

26. Fromentin, R. et al. CD4+ T Cells Expressing PD-1, TIGIT and LAG-3 Contribute to HIV Persistence during ART. PLoS Pathog 12, e1005761 (2016).

27. Descours, B. et al. CD32a is a marker of a CD4 T-cell HIV reservoir harbouring replication-competent proviruses. Nature 2017 543:7646 543, 564–567 (2017).

28. Osuna, C. E. et al. Evidence that CD32a does not mark the HIV-1 latent reservoir. Nature 561, E20–E28 (2018).

29. Veazey, R. S. et al. Dynamics of CCR5 expression by CD4(+) T cells in lymphoid tissues during simian immunodeficiency virus infection. J Virol 74, 11001–11007 (2000).

30. Chahroudi, A. et al. Target Cell Availability, Rather than Breast Milk Factors, Dictates Mother-to-Infant Transmission of SIV in Sooty Mangabeys and Rhesus Macaques. PLoS Pathog 10, e1003958 (2014).

31. Pandrea, I. et al. Mucosal simian immunodeficiency virus transmission in African green monkeys: susceptibility to infection is proportional to target cell availability at mucosal sites. J Virol 86, 4158–4168 (2012).

32. Xu, Y. et al. HIV-1 and SIV Predominantly Use CCR5 Expressed on a Precursor Population to Establish Infection in T Follicular Helper Cells. Front Immunol 8, (2017).

33. Contento, R. L. et al. CXCR4-CCR5: A couple modulating T cell functions. Proc Natl Acad Sci U S A 105, 10101–10106 (2008).

34. Zaunders, J. J. et al. Increased Turnover of CCR5+ and Redistribution of CCR5-CD4 T Lymphocytes during Primary Human Immunodeficiency Virus Type 1 Infection. J Infect Dis 183, 736–743 (2001).

35. Zaunders, J. J. et al. Early proliferation of CCR5+ CD38+++ antigen-specific CD4+ Th1 effector cells during primary HIV-1 infection. Blood 106, 1660–1667 (2005).

36. Zaunders, J. J. et al. CD127 + CCR5 + CD38 +++ CD4 + Th1 Effector Cells Are an Early Component of the Primary Immune Response to Vaccinia Virus and Precede Development of Interleukin-2 + Memory CD4 + T Cells. J Virol 80, 10151–10161 (2006).

37. Bleul, C. C., Wu, L., Hoxie, J. A., Springer, T. A. & Mackay, C. R. The HIV coreceptors CXCR4 and CCR5 are differentially expressed and regulated on human T lymphocytes. Proc Natl Acad Sci U S A 94, 1925–1930 (1997).

38. Zaunders, J. et al. Mapping the extent of heterogeneity of human CCR5R CD4R T cells in peripheral blood and lymph nodes. AIDS 34, 833–848 (2020).

39. Xu, Y. et al. HIV-1 and SIV predominantly use CCR5 expressed on a precursor population to establish infection in T follicular helper cells. Front Immunol 8, 376 (2017).

40. Brelot, A. & Chakrabarti, L. A. CCR5 Revisited: How Mechanisms of HIV Entry Govern AIDS Pathogenesis. J Mol Biol 430, 2557–2589 (2018).

41. Glass, W. G. et al. CCR5 deficiency increases risk of symptomatic West Nile virus infection. Journal of Experimental Medicine 203, 35–40 (2006).

42. Lim, J. K. et al. Genetic Deficiency of Chemokine Receptor CCR5 Is a Strong Risk Factor for Symptomatic West Nile Virus Infection: A Meta-Analysis of 4 Cohorts in the US Epidemic. J Infect Dis 197, 262–265 (2008).

43. Lim, J. K. et al. CCR5 Deficiency Is a Risk Factor for Early Clinical Manifestations of West Nile Virus Infection but not for Viral Transmission. J Infect Dis 201, 178–185 (2010).

## METHODS REFERENCES

1. William Chang, W. L. et al. RhcMV serostatus and vaccine adjuvant impact immunogenicity of RhcMV/SiV vaccines. Sci Rep 10, 14056 (2020).

2. Amedee, A. M. et al. Early Sites of Virus Replication after Oral SIV mac251 Infection of Infant Macaques: Implications for Pathogenesis. AIDS Res Hum Retroviruses 34, 286–299 (2018).

3. Merriam, D. et al. Depletion of Gut-Resident CCR5+ Cells for HIV Cure Strategies. AIDS Res Hum Retroviruses 33, S70–S80 (2017).

4. Schmitz, J. E. et al. Control of viremia in simian immunodeficiency virus infection by CD8+ lymphocytes. Science (1979) 283, 857–860 (1999).

5. Schmitz, J. E. et al. A nonhuman primate model for the selective elimination of CD8+ lymphocytes using a mouse-human chimeric monoclonal antibody. Am J Pathol 154, 1923–1932 (1999).

6. Okoye, A. A. et al. Early antiretroviral therapy limits SIV reservoir establishment to delay or prevent post-treatment viral rebound. Nat Med 24, 1430–1440 (2018).

7. Cline, A. N., Bess, J. W., Piatak, M. & Lifson, J. D. Highly sensitive SIV plasma viral load assay: practical considerations, realistic performance expectations, and application to reverse engineering of vaccines for AIDS. J Med Primatol 34, 303–312 (2005).

8. Deere, J. D. et al. SARS-CoV-2 Infection of Rhesus Macaques Treated Early with Human COVID-19 Convalescent Plasma. Microbiol Spectr 9, e0139721 (2021).

9. Ma, Z.-M., Dutra, J., Fritts, L. & Miller, C. J. Lymphatic Dissemination of Simian Immunodeficiency Virus after Penile Inoculation. J Virol 90, 4093–4104 (2016).

10. Schindelin, J. et al. Fiji: an open-source platform for biological-image analysis. Nat Methods 9, 676–682 (2012).

11. Hill, A. L. et al. Real-Time Predictions of Reservoir Size and Rebound Time during Antiretroviral Therapy Interruption Trials for HIV. PLoS Pathog 12, e1005535 (2016).

12. Hill, A. L., Rosenbloom, D. I. S., Fu, F., Nowak, M. A. & Siliciano, R. F. Predicting the outcomes of treatment to eradicate the latent reservoir for HIV-1. Proc Natl Acad Sci U S A 111, 13475–13480 (2014).

13. Rosenbloom, D. I. S. et al. Designing and Interpreting Limiting Dilution Assays: General Principles and Applications to the Latent Reservoir for Human Immunodeficiency Virus-1. Open Forum Infect Dis 2, ofv123 (2015).

